# Biological and synthetic surfactant exposure increase anti-microbial gene occurrence in a freshwater mixed microbial biofilm environment

**DOI:** 10.1101/2022.09.14.507961

**Authors:** Stephanie P. Gill, William J. Snelling, James S.G. Dooley, Nigel G. Ternan, Ibrahim M. Banat, Joerg Arnscheidt, William R. Hunter

## Abstract

Aquatic habitats are particularly susceptible to chemical pollution from domestic, agricultural, and industrial sources. Antimicrobials are commonly used in medical and industrial environments to reduce harmful bacteria and biofilms. This has led to the rapid increase in the prevalence of antimicrobial resistant (AMR) genes. Alternate remedies to fight pathogenic bacteria and biofilms are in development including synthetic and biological surfactants such as sodium dodecyl sulphate (SDS) and rhamnolipids respectively. In the aquatic environment these surfactants are present as pollutants with potential to affect biofilm formation and AMR gene occurrence; however, there is limited research showing the actual environmental impact of such exposure. We tested the effects of rhamnolipid and SDS on natural aquatic biofilms in a freshwater stream in Northern Ireland. We grew biofilms on contaminant exposure substrata deployed within the stream over four weeks, and then carried out shotgun sequencing to determine microbial community composition, through 16s rRNA analyses (64,678 classifiable reads identified), and AMR gene occurrence (81 instances of AMR genes over 9 AMR gene classes) through a metagenomic analysis. There were no significant changes in community composition within all systems; however, biofilm exposed to rhamnolipid had a greater number of unique taxa as compared to our SDS treatments and controls. AMR gene prevalence was higher in surfactant-treated biofilms, with biofilm exposed to rhamnolipids having the highest presence of AMR genes and classes compared to the control or SDS treatments, in which genes encoding for rifampin resistance were detected. Our results suggest that the presence of rhamnolipid, and to a lesser extent SDS, encourages an increase in the prevalence of AMR genes in biofilms produced in mixed use water bodies.

## Introduction

Chemical contamination from point and non-point source pollution poses a major problem for our environment; as industries and the human population expand more synthetic and anthropogenic toxic chemicals are released into the waterways from domestic, agricultural, and industrial sources (Yang *et al*. 2016, Fang *et al*. 2019). Chemical pollution encourages the increase of resistance gene selection in bacteria. The continued presence of antimicrobials, heavy metals, and biocidal chemicals leads to an increasingly common selection for resistance genes, as the need for bacteria to protect themselves against a chemical hazard outweighs any cost of carrying the gene (Singer *et al*. 2016). Antimicrobial chemicals are used in medical environments for surface and device disinfection. Unfortunately, overuse of these chemicals has led to an increase in bacterial resistance to them, with correlations showing increases in antimicrobial resistance (AMR) genes in areas with higher antimicrobial chemical presence (Maillard 2005, Hartmann *et al*. 2016). Based on metagenomic analyses AMR gene abundance is estimated to vary significantly across countries, with higher abundances more prevalent in areas with socio-economic disadvantages (Hendriksen *et al*. 2019). Accurate attribution of global deaths to AMR remains a challenge. Recent estimates are at an approximate annual death toll of 4.95 million people and extrapolation of available data suggests further increases for the future (Dunachie *et al*. 2020, Murray *et al*. 2022).

Many microorganisms, including various bacterial taxa, preferentially reside within biofilms where they are protected from a variety of environmental stressors (Bernbom *et al*. 2011, DePas *et al*. 2014, Rode *et al*. 2020). Biofilms provide an increased protection from antibiotics as the extracellular polymeric substance (EPS) matrix surrounding biofilms limits antibiotic penetration (De Beer *et al*. 1994, Mah and O’Toole 2001). Within this sheltered environment, horizontal gene transfer takes place which can increase the levels of AMR genes present within a biofilm in comparison to the outside. Environmental biofilms are also common carriers and habitats of pathogenic and opportunistic bacteria such as *Escherichia coli, Legionella* spp., and *Pseudomonas aeruginosa* (Wingedner and Flemming, 2011). Horizontal gene transfer of AMR genes can be increased by pathogenic bacteria residing within biofilms making them potential hotspots of AMR genes in the right conditions (Bowler *et al*. 2020, Ma *et al*. 2021).

Biosurfactants, or naturally produced surfactants (synthetic chemicals that alter surface tension between liquids), play a major role at the early development stage of biofilms mainly through the maintenance of channels through the matrix which enhance the exchange of nutrients and gases with the ambient environment. Ultimately biosurfactants also result in the dissociation of biofilm surface layers into planktonic mobile forms (Marchant and Banat 2012, Quinn *et al*. 2013; Banat *et al*. 2014). Although biosurfactants can naturally control biofilm development, and to some extent their structure, once too large, higher concentrations can be used to irradicate or inhibit biofilms (Satpute *et al*. 2016). In industrial and medical settings synthetic surfactants are typically used to remove unwanted biofilms; however, they also have found a large and increasing domestic use as essential ingredients in shampoos, soaps, toothpastes, and detergents (Ivankovic and Hrenovic 2010). For a variety of medical and technical applications the use of surfactants, both alone and in combination with antibiotics, has been proposed to break down harmful bacterial biofilms thereby reducing the reservoir of AMR genes (Zhang *et al*. 2016, Wang *et al*. 2020). However, growing awareness of the synthetic surfactants’ environmental toxicity has focused more effort on the investigation of safer alternatives, such as biosurfactants (Paraszkiewicz *et al*. 2021). One of the candidate substances in such tests is the biosurfactant rhamnolipid which in combination with antibiotics was able to remove 95% of tested biofilms after exposure (Chen *et al*. 2019).

Research concerning the effects of surfactants and biosurfactants on biofilms and changes in abundance and diversity of AMR genes has hitherto only concerned industrial and medical biofilms, while environmental settings remained underexposed. Therefore, this study investigated how a biological surfactant, rhamnolipid, and a synthetic surfactant, SDS, affected the development of microbial biofilms in a mixed land-use stream environment. We aimed to determine how exposure to both surfactants influenced microbial community composition and AMR gene prevalence of the exposed biofilms. We hypothesized that in a natural aquatic ecosystem, rhamnolipid and SDS exposure would alter biofilm community composition to favor taxa less sensitive to chemical exposure while also decreasing AMR gene abundance. Community composition changes were quantified through a 16s rRNA analysis. AMR gene abundances were examined through a metagenomic analysis of resistance to elfamycin, amingoclycoside, tetracycline, isoniazid, aminocoumarin, fluroquinolone, and rifampin.

## Experimental Procedures

### Experimental set up

The Ballysally Blagh is a second-order lowland stream at the town of Coleraine near the North Coast of Northern Ireland, UK. This tributary to the lower River Bann has an average mainstream channel slope (S1085) of 6.22 m km^-1^ between points of 10% and 85% of mainstream length above catchment outlet (Gardner and Wilcock, 2003). It drains a 14.2 km^2^ catchment of water gley soils with mixed land cover, i.e. grassland 55.9%, arable/horticultural 21.9%, bog 13.7%, urban 7.3%, woodland <2% (National River Flow Archive 2022). The non-urban area is predominantly used for agricultural purposes (McClean and Hunter 2020). The Ballysally Blagh’s baseflow index of 0.51 characterizes an intermediate level of water transfers towards its stream flow from storage in superficial deposits of mixed permeability. Artificial drainage and stream channel profiling have resulted in a flashy stream discharge response to precipitation as indicated by the stream’s Q_5_:Q_95_ discharge ratio of 24 (National River Flow Archive station 203050).

Nine contaminant exposure substrata (CES) cups were prepared as per Costello *et al*. (2015) using glass fiber (GF/C) filters. Clean tea filters (Lipton polypropylene nonwoven mesh (<0.25 mm) (Mori et al. 2021)) were added into each lid as a particulate sieve to minimize sedimentation on the glass fiber filters. Three cups contained agarose gel dosed with 150 ppm rhamnolipid (JBR 425, Jeneil Biosurfactant Company, Wisconsin USA), three cups contained agarose gel dosed with an equimolar concentration of SDS (75 ppm) (BDH, AnalaR, Poole, England) and the final three cups were controls with agarose gel only. Cups were zip tied together and attached to a large, submerged metal bar to secure their position on the stream bed of the Ballysally Blagh at 55°08’44.4”N 6°40’19.0”W.

### Sample Collection

CES cups were deployed in the stream for a period of four weeks from February 2020 to March 2020. To account for air contamination, three additional GF/C filters were used when CES cups were collected and exposed to air bacteria at the collection site. These three filters were then treated identically to all other filters from the experiment for the rest of the study and allowed the identification of any potential external microbial contaminants that were not from the stream environment. All cups were rinsed in ultrapure water to remove any debris that collected on top of each cup and filter. Individual sterile tweezers were then used to remove the GF/C filters with developed biofilm from the CES cups under a sterile laminar flow hood. Filters were placed immediately into sterile test tubes, sealed, and frozen at -20°C.

DNA extractions were performed using a FastDNA Spin kit for Soil (MP Biomedical, Leicester, UK) as per the manufacturer’s guidelines. Biofilm material was scraped off the filters recovered from the stream before performing DNA extractions. In the absence of visible biofilm on the air controls the actual filters were processed to obtain any DNA that may have originated from contamination. To confirm DNA presence within each sample, DNA was quantitated using a ND1000 Nanodrop Spectrophotometer (Labtech Int., Heathfield, UK). Shotgun sequencing was performed with a MinION portable sequencer and R9 flow cell (Oxford Nanopore, Oxford, UK), in accordance with manufacturer’s guidelines, including the Agencourt AMPure XP Bead (Beckman-Coulter, High Wycombe, UK) clean-up step, using a Rapid Barcoding Sequencing kit SQK-RBK004 (Oxford Nanopore, Oxford, UK) with a minimal acceptable DNA quality score of 7. Similar DNA yields were used for sequencing to allow for appropriate comparison between samples, with each of the twelve samples analyzed in triplicates that were then averaged. All samples from the main experiment, with the exception of one low yield experimental control, yielded 280 ng of DNA. Air controls and the single low yield experimental control yielded 48 ng of DNA. Results from the low yield experimental control, and the high yield experimental control samples were compared before being averaged later in the analysis protocol to ensure similar species abundance ratios occurred.

### Data analysis

The twelve resultant Fast Q files from sequencing were processed individually using the Fastq WIMP (v2021.03.05) for utilizing the 16s rRNA workflow to obtain bacterial species identification and Fastq Antimicrobial Resistance (v2021.05.17) for identification of AMR gene prevalence as well as changes to efflux pumps conferring antibiotic resistance, and peptide antibiotic resistance genes software (Oxford Nanopore). Recommended settings were used along with a minimum DNA quality score of 7 to obtain higher quality sequences, where all 326,805 sequences used had an average quality score of 10. Results from both software packages were downloaded as CSV files for further analysis in R statistical software. Any bacterial species in the air control samples identified with the Fastq WIMP 16s rRNA workflow were removed from all files before further analyses continued based on the assumption that these were contaminating species of airborne bacteria.

Statistical analysis employed R version 4.0.3 (R Core Team 2020) and the R packages car for normality testing (Fox and Weisberg, 2018), vegan for univariate and multivariate statistics (Oksanen *et al*. 2020), and ggplot2 for figure creation (Wickham, 2016). Past 3 software (Hammer *et al*. 2001) was used to perform all non-metric multidimensional scaling (NMDS) analyses. All univariate data met assumptions of normality and homogeneity of variance.

Significance was assumed at *p*< 0.05. Diversity was first examined in the 16s rRNA results using the Shannon-Wiener diversity index (examining abundance and evenness of species present). We included all singletons and doubletons in this to account for potentially rare species. Diversity index values were analyzed with a one-way analysis of variance (ANOVA) test to determine if any diversities were significantly different from each other (*p*< 0.05). Singletons and doubletons were then removed, and data were rarified and subsampled. This action allowed us to account for the differences in DNA yields within our data by keeping the same ratio of organisms within each sample but reducing abundance where there was an equal amount of total taxa in each sample. An NMDS was then performed followed by an analysis of similarities (ANOSIM) to test for significant differences among the community composition of biofilms.

AMR gene presence, i.e. frequency of presence or absence of each gene as noted by the Fastq Antimicrobial Resistance software, was examined first with an NMDS and then a permutational multivariant analysis of variance (PERMANOVA). This determined if there were differences among both frequency of presence of detected AMR genes and the types of gene. We then ran an ANOVA to identify significant differences in only the types of genes present.

## Results

### Community composition

First, we determined how the community composition of biofilms shifted after exposure to both SDS and rhamnolipids. Overall, we found large dominance by *Escherichia, Acidovorax, Janthinobacterium, Shigella, Flavobacterium*, and *Pseudomonas* spp. in all treatments and controls. The rhamnolipid treated samples contained 566 unique taxa, SDS treatments had 478, while control treatments had the lowest at 386. However, despite having the lowest total taxonomic richness, the controls still had 57 unique taxa when compared to the treatments. Rhamnolipid had 177 unique taxa and SDS had 96.

We found no significant differences in microbial diversity among any of the biofilms (ANOVA F=1.409, p=0.315) (Figure 1A). Although biofilm in SDS treatments had a higher taxonomic diversity than in rhamnolipid treatments, the control samples had the largest range of diversity, accounting for both the highest and lowest diversity overall. Consequently, while there were some differences in the diversity arising in SDS and rhamnolipid treatments, neither treatment appeared significantly different to the control. The employed diversity measurements examine both abundance and evenness, implying that although rhamnolipid treatments contained a greater number of unique taxa, most likely that many of these were low abundance taxa which reduced the treatment’s overall diversity score.

**Figure 1.**
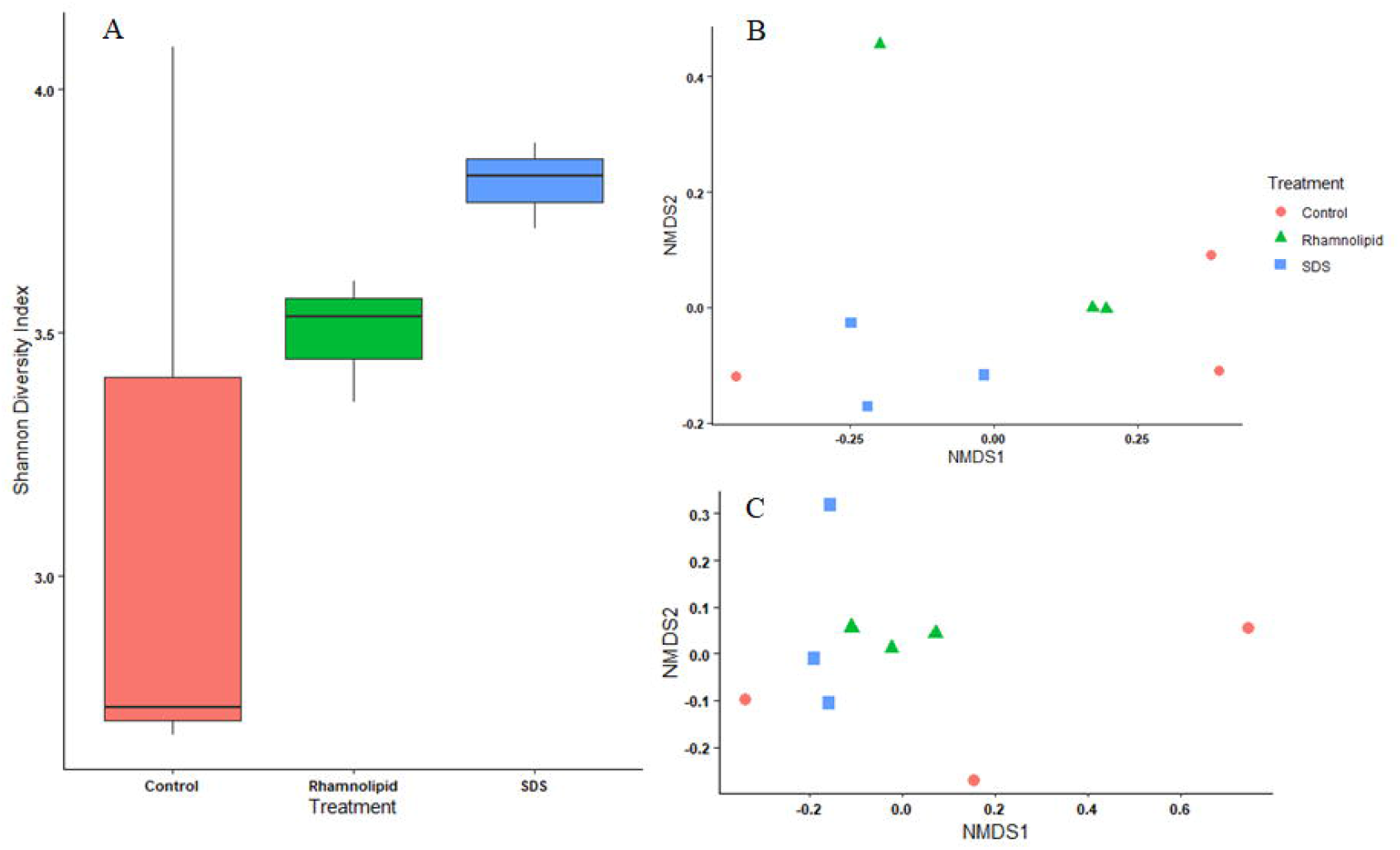
Microbial diversity with the (A) Shannon Wiener diversity index and two NMDS analyses depicting community differences of naturally grown microbial biofilms continuously exposed to 150 ppm of the biosurfactant rhamnolipid and 75 ppm of the synthetic surfactant sodium dodecyl sulphate (SDS) over a period of 4 weeks in a stream with a mixed land use catchment (B) with and (C) without the top ten most dominant genera identified. These mixed biofilms were naturally developed using contaminant exposure substrate cups.

NMDS was used to identify differences in the overall community composition of treatment groups, which would demonstrate if exposure to either chemical had altered biofilm composition. There were no distinct groupings within the NMDS (Figure 1B), which was confirmed by an ANOSIM (R=0.1512, *p*= 0.226), implying a lack of unique community compositions within the biofilms. To identify if there were differences among the rarer taxa within our samples, the NMDS (Figure 1C) and ANOSIM (R=0.1111, *p*= 0.14) were rerun without the top ten most dominant taxa and no significant differences or groupings were identified within the rarer taxa across the treatments and controls.

Dominant taxa were also similar among treatments. All treatments and controls were dominated by *Escherichia, Acidovorax, Janthinobacterium, Shigella, Flavobacterium*, and *Pseudomonas* spp. With the exception of *Ralstonia* and *Leptothrix*, where the former was only present in the control, and the latter only present in the rhamnolipid and SDS treatments, lists of the top ten genera identified within each treatment and control were almost identical (Table 1). While the former two genera were not among the top ten for all groups, they still present in each control or treatment albeit sometimes at lower abundances. Although there was considerable overlap in dominant genera within each treatment, abundance of taxa identified (based on 16s rRNA sequences) within each genera differed (Figure 2). Rhamnolipid treatments had the greatest concentration of *Escherichia*, while the control had the greatest concentrations of *Shigella*, and SDS had the greatest concentrations of *Janthinobacterium*. For several genera such as *Escherichia, Janthinobacterium*, and *Shigella* the control and rhamnolipid treatments had more comparable abundances than the rhamnolipid and SDS treatments. Rhamnolipid and SDS treatments had similar *Pseudomonas* concentrations when compared to the control. Therefore, it is reasonable to infer that each genus had its individual sensitivity, and *Shigella* spp. may be the most sensitive to chemical contamination as compared to *Escherichia* or *Janthinobacterium*.

**Table 1.**
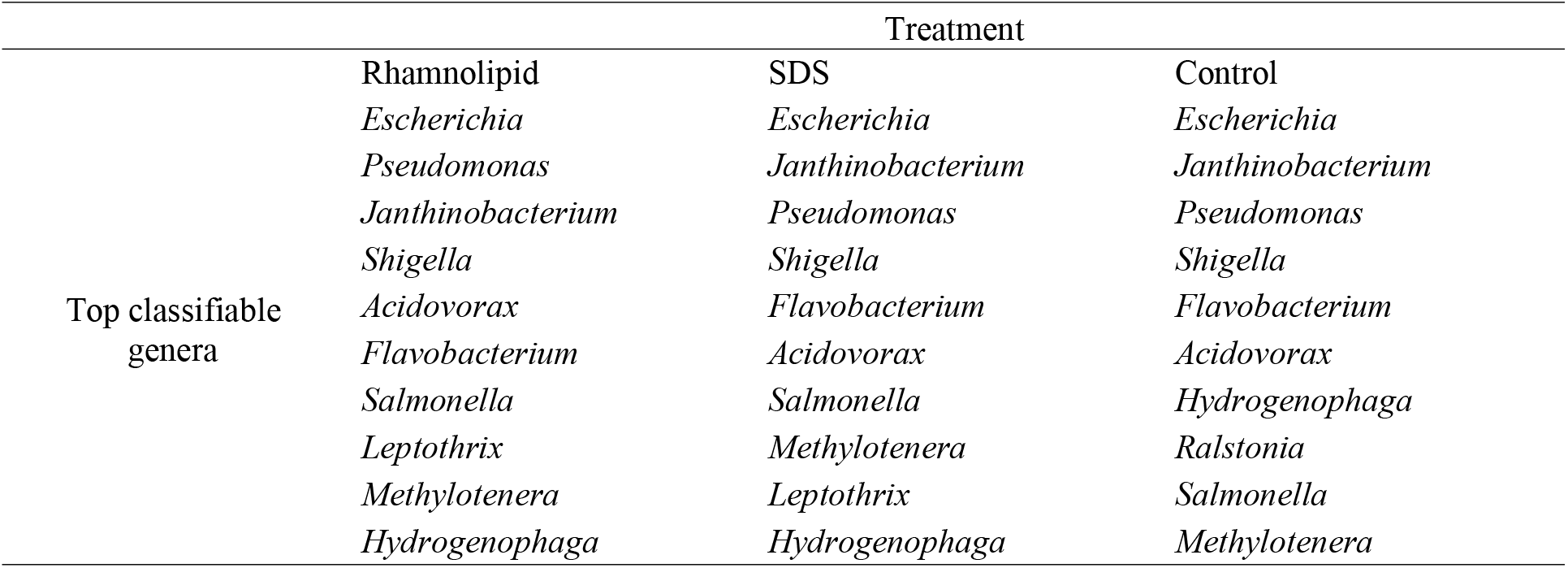
Top 10 classifiable genera within microbial biofilms continuously exposed to 150 ppm of the biosurfactant rhamnolipid and 75 ppm of the synthetic surfactant sodium dodecyl sulphate (SDS) over a period of 4 weeks in a stream. Biofilms were mixed and naturally developed using contaminant exposure substrata cups. Genera are listed in order of dominance within each treatment.

**Figure 2.**
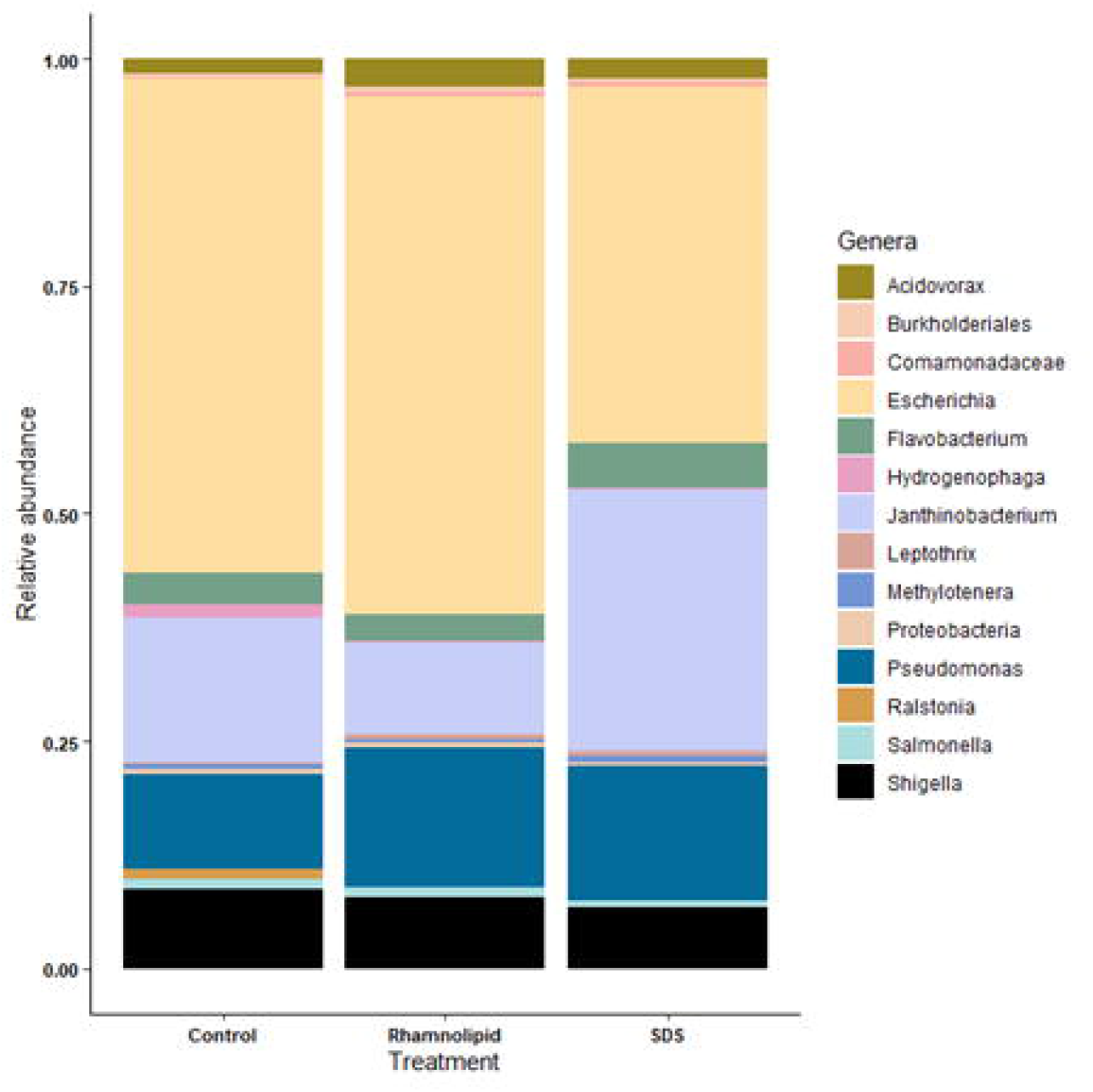
Relative abundances of the most dominant genera, as measured by sequence abundance, within microbial biofilm communities from mixed biofilms grown in contaminant exposure substrate cups and continuously exposed to 150 ppm of the biosurfactant rhamnolipid and 75 ppm of the synthetic surfactant sodium dodecyl sulphate (SDS) over a period of four weeks in a stream with a mixed land use catchment.

### Antimicrobial Resistance Genes

There were no statistically significant differences in abundance of AMR genes within each treatment (PERMANOVA F=1.3812, *p*=0.195). This was confirmed by NMDS as there were no distinct groupings among treatments and controls (Figure 3). However, further examination of the differences in AMR genes showed generally higher numbers of AMR genes within the rhamnolipid treated cups when compared to the SDS treatments and the controls (Figure 4A), although the difference was not significant (ANOVA F=1.926, *p*=0.226). The presence of each AMR gene was also not significantly different across treatments (MANOVA: elfamycin F=3, *p*=0.125; aminoglycoside F=1.1, *p*=0.392; tetracycline F=1.78, *p*=0.113; isoniazid F=1, *p*=0.422; efflux pumps F=0.676, *p*=0.544; aminocoumarin F=0.5, *p*=0.629; fluoroquinolone F=1.75, *p*=0.252; rifampin F=1, *p*=0.422; peptide F=0.5, *p*=0.629). Overall 81 instances of AMR gene presence were identified across 9 AMR classes. The rhamnolipid treatment was the only treatment in which genes encoding elfamycin, isoniazid, and rifampin resistance were detected, and also had the highest abundance of aminoglycoside resistance genes. Tetracycline and aminocoumarin resistance was also only detected in treatments, but not in the controls (Figure 4B).

**Figure 3.**
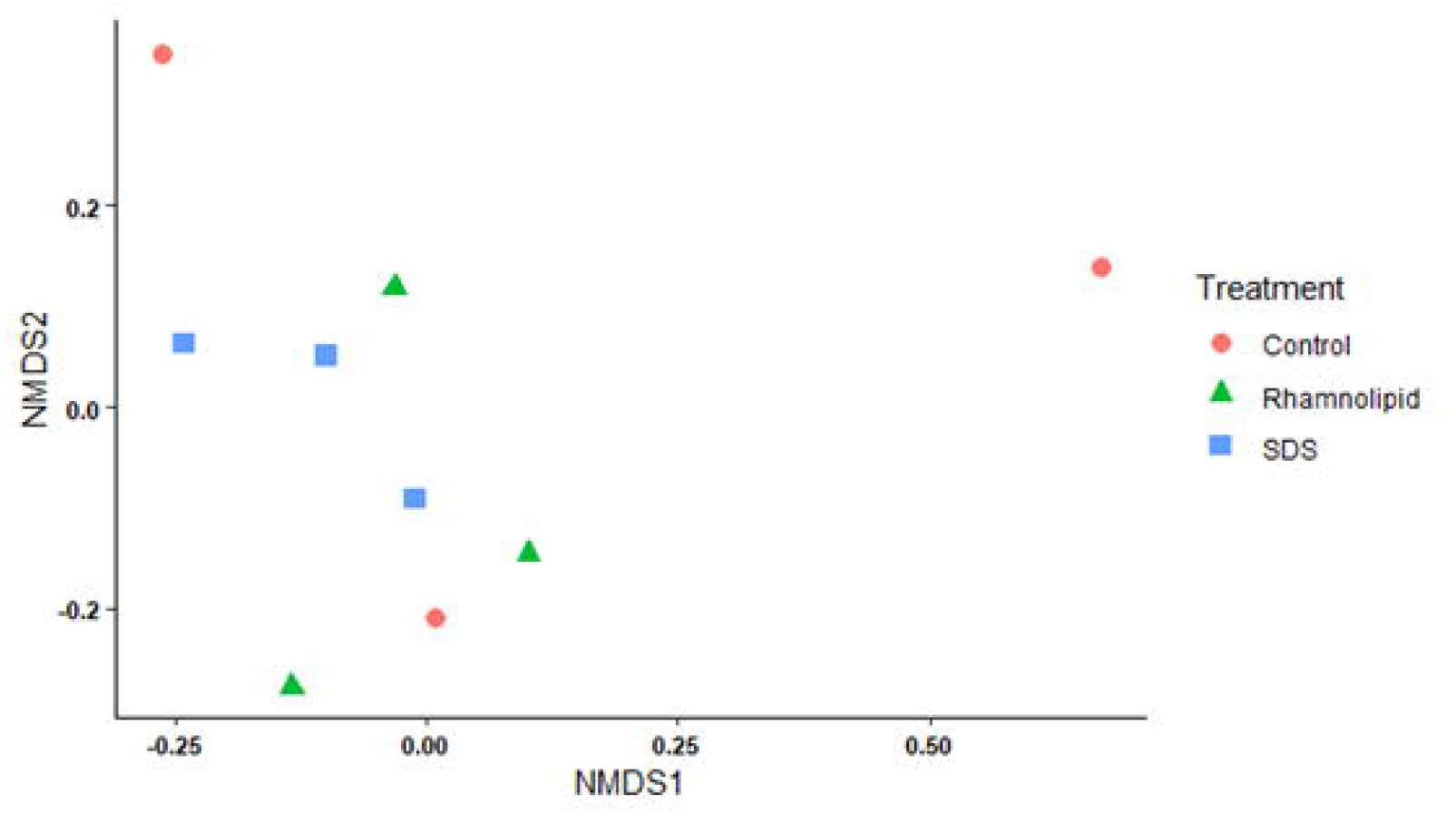
NMDS depicting the differences in antimicrobial resistance genes identified within mixed microbial biofilms grown in contaminant exposure substrata cups and continuously exposed to 150 ppm of the biosurfactant rhamnolipid and 75 ppm of the synthetic surfactant sodium dodecyl sulphate (SDS) over a period of 4 weeks in a stream with a mixed land use catchment.

**Figure 4.**
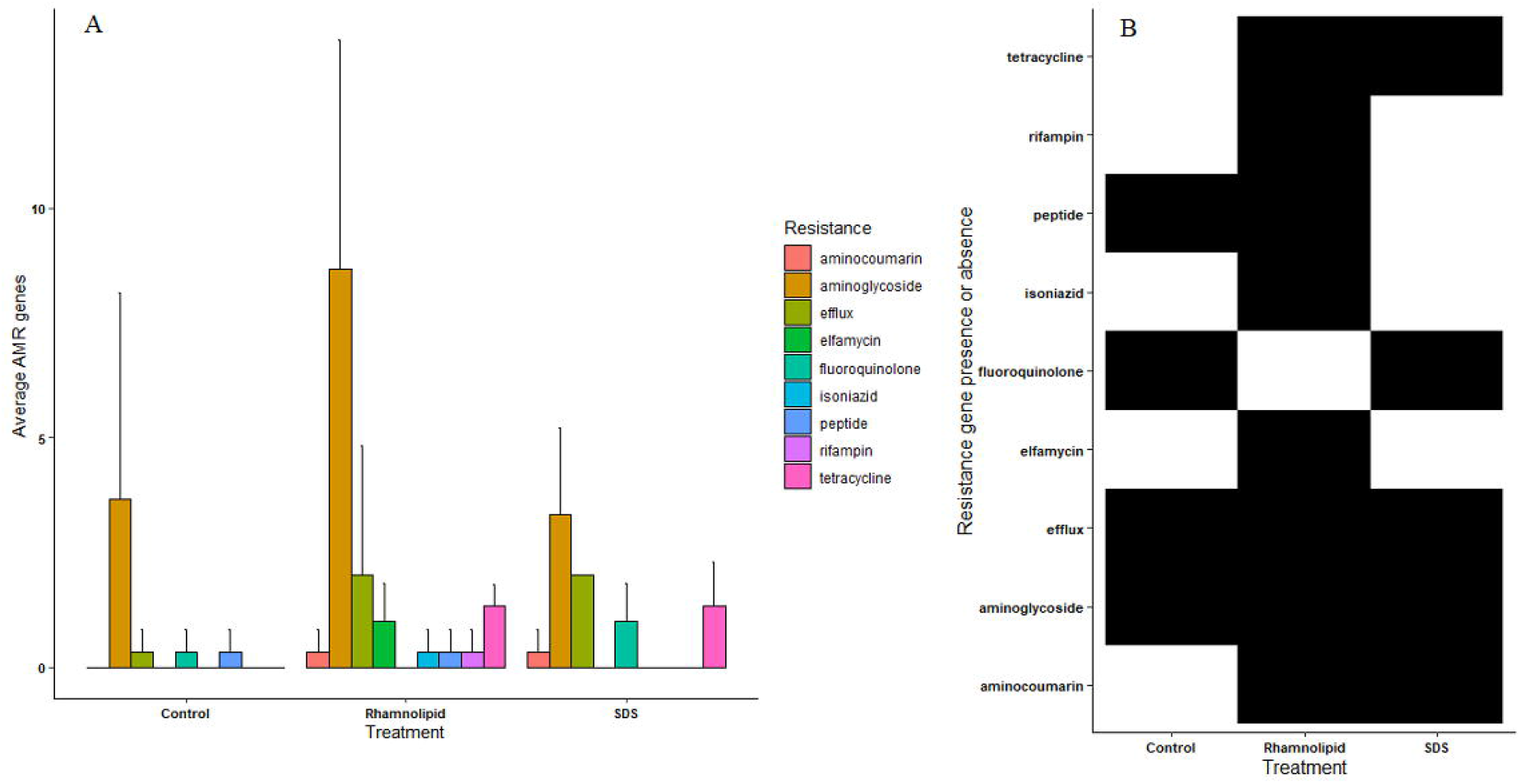
Differences in (A) average number of antimicrobial resistance (AMR) genes present and (B) the type of AMR genes present, where black signifies presence and white signifies absence, within a mixed microbial biofilm grown in contaminant exposure substrate cups continuously exposed to 150 ppm of the biosurfactant rhamnolipid and 75 ppm of the synthetic surfactant sodium dodecyl sulphate (SDS) over a period of 4 weeks in a stream with a mixed land use catchment.

## Discussion

This study aimed to determine how rhamnolipid and SDS exposure affects the occurrence of AMR genes in natural mixed biofilms. Although application of rhamnolipid and SDS treatments did not affect overall biofilm diversity and community composition, there were major differences in AMR gene prevalence. The unique presence of AMR genes was only identified after exposure to rhamnolipid with elfamycin, isoniazid, and rifampin resistance. Tetracycline and aminocoumarin resistances were also identified in biofilms after exposure to either surfactant which is troubling as surfactants are used to enhance antibiotic potency.

The Ballysally Blagh drains a catchment that is 92.7 % agricultural and 7.3 % urban land (McClean and Hunter 2020, Hunter *et al*. 2021). As such, it regularly receives significant inputs of fecal material as a consequence of agricultural slurry spreading, and from a small sewage treatment works located upstream of the university campus. A wet February (right after closed season ends for slurry application) would also imply that extra fecal material from initial slurry application on waterlogged soil and wastewater from sewage overflow was present in the waterways. The Ballysally Blagh’s microbial community was dominated by *Escherichia, Acidovorax, Janthinobacterium, Shigella, Flavobacterium*, and *Pseudomonas* spp. within it, along with several other similar genera. *Escherichia* (Carlos *et al*., 2010) and *Shigella* (Olalemi and Dauda 2018) are commonly used as markers of fecal contamination in waterways from both human and animal sources. *Pseudomonas* and *Salmonella* bacterial presence are also used as an alternate indicator of fecal contamination due to their high presence in contaminated waters (Liang *et al*. 2015, Januário *et al*. 2020). All dominant genera are found in agricultural and urban environments (Cabral 2010, Rosi *et al*. 2018), with some of them already having developed resistance to antibiotic and chemical exposure (Kumar *et al*. 2019, Pang *et al*. 2019, Saraceno *et al*. 2021). *Pseudomonas* and *Acidovorax* spp. are extremely adept at survival in polluted environments, dominating once diverse habitats after chlorine exposure (Jia *et al*. 2015).

The lack of differences identified in the community composition between treatments is likely to be caused by the domination of these bacterial taxa, probably due to the stream being highly exposed to agricultural and urban land. Over 1/3 of the present taxa within our treatments and controls were made up of the same genera, and only two genera *Ralstonia* and *Leptothrix* were found in unique treatments. The greater dominance of *Ralstonia*, a potential plant pathogen (Wei *et al*. 2018), within the control treatment may indicate greater sensitivity to chemical exposure. Alternatively, the greater concentrations of *Leptothrix* within the surfactant treatments may indicate lower chemical sensitivity. *Escherichia* growth was also fairly limited within SDS treated systems when compared to the control and rhamnolipid treatments and *Janthinobacterium* growth was increased, potentially indicating less sensitivity of *Janthinobacterium* to SDS.

More taxa were identified within the rhamnolipid and SDS treatments as compared to the control. Typically, exposure to either chemical has led to a reduced taxa count (Gill *et al*. 2022). In our experiments, exposure to rhamnolipids yielded the highest number of unique taxa, SDS the second highest, and control the lowest, following the pattern of AMR gene abundance and diversity as well. Unique taxa included *Stella*, a star shaped bacterium commonly found in sewage and soil/freshwater environments (Vasilyeva 1985) which was observed in the rhamnolipid treatment only, and *Haliangium*, a bacterial genus previously found in wastewater treatment plant sludge (Gao *et al*., 2016) which was observed in the SDS treatments only. Although the taxa were unique to their treatments, they are commonly found in environments exposed to sewage and such conditions tend to facilitate horizontal gene transfer and increase the abundance of AMR genes (Uluseker *et al*., 2021). We propose the possibility that surfactants may enhance the passage of AMR genes into bacterial cells. Surfactants, such as SDS, can disrupt cell membranes (Islam *et al*., 2017), potentially making uptake of genes from the environment, possibly through horizontal gene transfer, easier. However, we also emphasize that as there were taxa unique to the control samples, the presence of either surfactant may also make a habitat more inhospitable for some taxa.

Treatments and controls all acquired AMR genes during the course of the experiment, which is to be expected considering the high levels of resistance typically found in *Escherichia* and *Pseudomonas* spp. (Tadesse *et al*. 2012, Pang *et al*. 2019) and the high likelihood of fecal pollution in our agricultural and urban stream. Previous research has identified *Escherichia* and *Pseudomonas* resistance to aminoglycoside (Garneau-Tsodikova and Labby 2016), fluoroquinolone (Redgrave *et al*. 2014), and tetracycline (Gasparrini *et al*. 2020) among others. However, rhamnolipid treated systems developed a greater abundance of AMR genes when compared to the control or SDS treatments, and they were also the systems with the highest concentrations of *Escherichia*. This implies that they may be able to promote the retention of AMR genes within a biofilm and potentially encourage growth of pathogenic strains of bacterial genera such as *Escherichia*. They were also the only treatments to affect AMR gene prevalence with *Streptomyces* (which had high prevalence of elfamycin resistance), and *Mycobacterium* (which had high prevalence of isoniazid resistance), and the only treatment to develop resistance to rifampin. This is problematic as rhamnolipids are currently researched for their ability to enhance antibiotic effects and antibiotic potency (Bharali *et al*. 2013, Chen *et al*. 2019).

Surfactants in medical (Deo *et al*. 2010) and industrial applications (Schlüter *et al*. 2007) have also been investigated for their ability to reduce the number of antibiotic resistant bacteria in hospital and care environments and in wastewater treatment plants. However, it has been noted that bacteria have started to develop resistance to surfactants, specifically quaternary ammonium compounds (QACs), and that bacterial strains with QAC resistance are more likely to co-select for antibiotic resistance as well (Gaze *et al*. 2005). While SDS and rhamnolipids are not QACs, our dataset suggests that there is a larger abundance and diversity of AMR genes developing among biofilm able to grow on an exposed substrate. Our results appear to be the first evidence to suggest that rhamnolipids may have a large impact on the abundance of AMR genes in the wild, and that SDS may also slightly encourage AMR gene selection. It is likely that although biosurfactants and surfactants may facilitate access of antibiotics to bacterial cells through weakening cell membranes, the beneficial action of combined treatments also leads to higher increases in AMR gene selection among surviving bacteria.

We sought to identify the effects of a surfactant and biosurfactant on the community composition and AMR gene abundance of a mixed microbial biofilm grown in a stream exposed to agricultural and urban sources of pollution. Based on previous research we assumed that biofilm composition would alter after exposure to rhamnolipid and SDS, and that there would be fewer AMR genes present (Zhang *et al*. 2016, Chen *et al*. 2019, Wang *et al*. 2020). However, we observed minimal compositional and diversity changes occurring, and an increase in AMR gene abundance after exposure. It is likely that an environment with a substantial presence of *Escherichia, Acidovorax, Janthinobacterium, Shigella, Flavobacterium*, and *Pseudomonas* favored opportunistic bacteria, allowing for biofilm compositional dominance by just a few species. These opportunistic bacteria are also well known for their selection of AMR genes in the presence of antibiotics and antimicrobial chemicals, which may have been encouraged by the presence of rhamnolipid and to a lesser extent of SDS. This would be extremely unfortunate as rhamnolipids are currently being studied for their ability to increase antibiotic potency in controlled experiments (Juma et al. 2020).

Our study is therefore the first example of this biosurfactant increasing AMR gene abundance, instead of acting as a biofilm control. Unless further field experiments are performed to supplement experiments under controlled laboratory conditions, it will be difficult to determine the impact of introducing rhamnolipids to agricultural and urban waterways. If they consistently increased abundance and diversity of AMR genes in natural biofilms then their presence in the aquatic ecosystem could be detrimental to both human and animal health as they could facilitate bacterial acquisition of multi-resistance against antibiotics, thus encouraging the evolution of a new superbug.

## Acknowledgements

The authors acknowledge the support of Ryan Lunenberg for assistance with microcosm placement. We also acknowledge Jeneil Biosurfactant, Saukville, WI, USA for providing rhamnolipid samples and the UK National River Flow Archive for the use of hydrological data for the Ballysally Blagh.

